# Nepicastat, a dopamine-beta-hydroxylase inhibitor decreases blood pressure and induces the infiltration of macrophages and B cells in the heart of spontaneous hypertensive rats

**DOI:** 10.1101/2023.07.28.549206

**Authors:** Shivanshu Chandan, Ganesh Kosher

**Affiliations:** Laboratory of Pharmacological Sciences, The Central University of Haryana, Haryana, India 123031

**Keywords:** Hypertension, Inflammation, Heart, Blood pressure

## Abstract

Nepicastat is a potent dopamine-beta-hydroxylase inhibitor that modulates the sympathetic nervous system by inhibiting the synthesis of norepinephrine. Nepicastat is a potential drug for the treatment of congestive heart failure. We sought to investigate the mechanistic role of Nepicastsat in the heart of Spontaneous Hypertensive Rats (SHR) rats. Here, we investigated if Nepicastat at both acute (7 days) and chronic administration (14 days) decrease blood pressure and echocardiography parameters in SHR rats. SHR 3-4 months male rats were administered either Nepicastsat (30mg/kg, orally), Enalapril (10 mg/kg, orally), or vehicle for 7 days or 14 days. Blood pressure and echocardiography parameters were recorded on day 0, day 3, day 7, and day 14 of drug administration. The animals were sacrificed, and tissues are collected for histology, qRTPCR, and flow cytometry analysis. At both acute and chronic administration, Nepicastat decreased systolic blood pressure and intraventricular septal thickness of SHR rats compared to vehicle groups. The decrease in blood pressure was comparable to Enalapril treated rats. Interestingly, Nepicastat also decreased the infiltrating macrophages and B cells in the hearts of SHR rats. In conclusion, Nepicastsat consistently decreased the systolic blood pressure but increased the macrophages and B cell infiltration in the heart of SHR rats.

## Introduction

Hypertension is one of the most important risk factors for several cardiovascular diseases (CVDs) including heart failure, and coronary heart disease^1-6^. Several antihypertensive drugs including angiotensin-converting enzyme inhibitors (ACEi), angiotensin II receptor blockers (ARB), calcium channel blockers (CCB), beta-blockers, and diuretics are currently used for the treatment of hypertension^7-9^. A combination of antihypertensive drugs is in use to keep in mind the multiple targets these therapies achieve. Despite these advanced therapeutic strategies, there is an urgent need for a new class of drugs in the last several years which can lower blood pressure with fewer side effects. Dopamine-β-hydroxylase (DBH), an enzyme of the sympathetic nervous system, is one such target system that plays a central role in regulating blood pressure^10-12^. DBH, catalyze the conversion of dopamine into noradrenaline thus balancing the vasoconstricting noradrenaline and vasodilating dopamine levels and ultimately regulating the blood pressure^13^. Owing to these functions, DBH inhibitors have been implicated in the treatment of hypertension and cardiac heart failure. DBH inhibitors gradually brings down the blood pressure instead of acute inhibition of blood pressure thus preventing the off-target effects of diuresis and natriuresis^14-16^. DBH inhibitors thus could be better alternatives to conventional drugs including beta blockers, ACE inhibitors, ARBs. A relatively few DBH inhibitors with moderate efficacies are available and none of them are used therapeutically^16^.

Among DBH inhibitors, nepicastat, etamicastat, and zamicastat are the most promising drugs but failed to reach clinical research owing to their side effects^15^. Nepicastsat is a DBH inhibitor that reduces blood pressure but crosses blood-brain barrier with the potential for neurological side effects^19^. Here, we investigated if acute or chronic administration of nepicastat lowers blood pressure with the potential off-target effects on other organs. We also compared the blood pressure-lowering effects of nepicastat with the conventional drug Enalapril. At both acute and chronic administration, Nepicastat decreased systolic blood pressure and intraventricular septal thickness of SHR rats compared to vehicle groups. The decrease in blood pressure was comparable to Enalapril treated rats. Interestingly, Nepicastat however increased the infiltration of macrophages and B cells in the hearts of SHR rats. In conclusion, Nepicastsat consistently decreased the systolic blood pressure and induced macrophage and B cell infiltration in the heart of SHR rats.

## Results

Baseline measurements of blood pressure revealed hypertension (≥ 180 mmHg) in all the animals. ECG parameters also suggested a decreased PR and QT interval in all the animals. Echo parameters revealed an increase in IVSd, IVSs, LVPWs, and LVPWd parameters in hypertensive animals. LVIDd and LVIDs parameters were decreased and thus suggested left ventricular hypertrophy in these animals. There was no significant difference in ejection fraction and fractional shortening parameters.

In a pilot study, we observed a decreased systolic blood pressure at 2 hrs. after drug (nepicastat and enalapril) administration. Enalapril administered SHR had reduced blood pressure to < 140 mmHg at 2 hrs. and even maintained till 24 after drug treatment. Nepicastat treatment however reduced the blood pressure (< 150mmHg) from 2 Hrs. – 12 hrs. but did not sustain the decreased blood pressure until 24 hrs. after drug administration (Fig. 1A).

**Figure 1.**
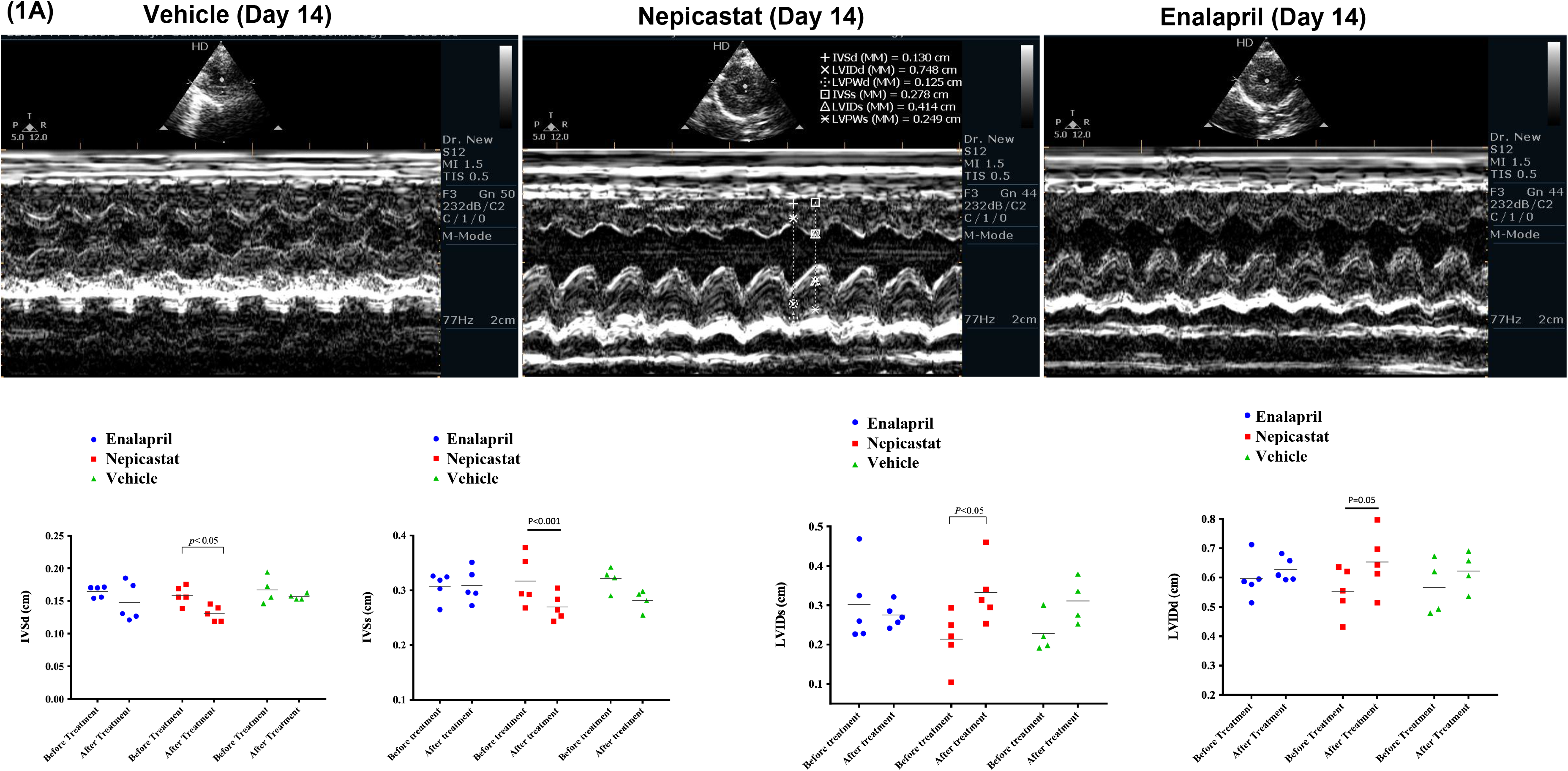

In the 5-days acute study,

We observed a significant (p<0.001 and p<0.0001) decreased BP in both nepicastat and enalapril administered rats when compared with baseline BP and vehicle group (Fig. 1B).

In a 10-day chronic study,

We observed a significant (p<0.05 and p<0.01) decreased BP at 6 hrs. in both nepicastat and enalapril administered rats when compared with baseline BP and vehicle group. We also observed a significant (p<0.001 and p<0.001) decrease in blood pressure at 24 hrs. in both nepicastat and enalapril administered rats when compared with baseline BP and vehicle group (Fig. 1C).

We did not find a significant difference in echo parameters IVSd, IVSs, LVIDd, LVIDs, and LVEF in rats administered with nepicastat when compared with the vehicle or enalapril group (Fig. 2A). Trichrome staining revealed a mild to moderate degree of fibrosis in all the SHR rats administered with either nepicastat, vehicle, or enalapril. There was decreased fibrosis or collagen deposition in nepicastat and enalapril administered rats when compared with the vehicle group (Fig. 2B). In addition, we did not find significant pathological changes in different tissues of rats administered with nepicastat. Histology however revealed a mild to moderate infiltration of inflammatory cells in the heart, lung, and kidney tissues of all the rats administered with either nepicastat, enalapril compared to vehicle-administered rats (Fig. 2C).

**Figure 2.**
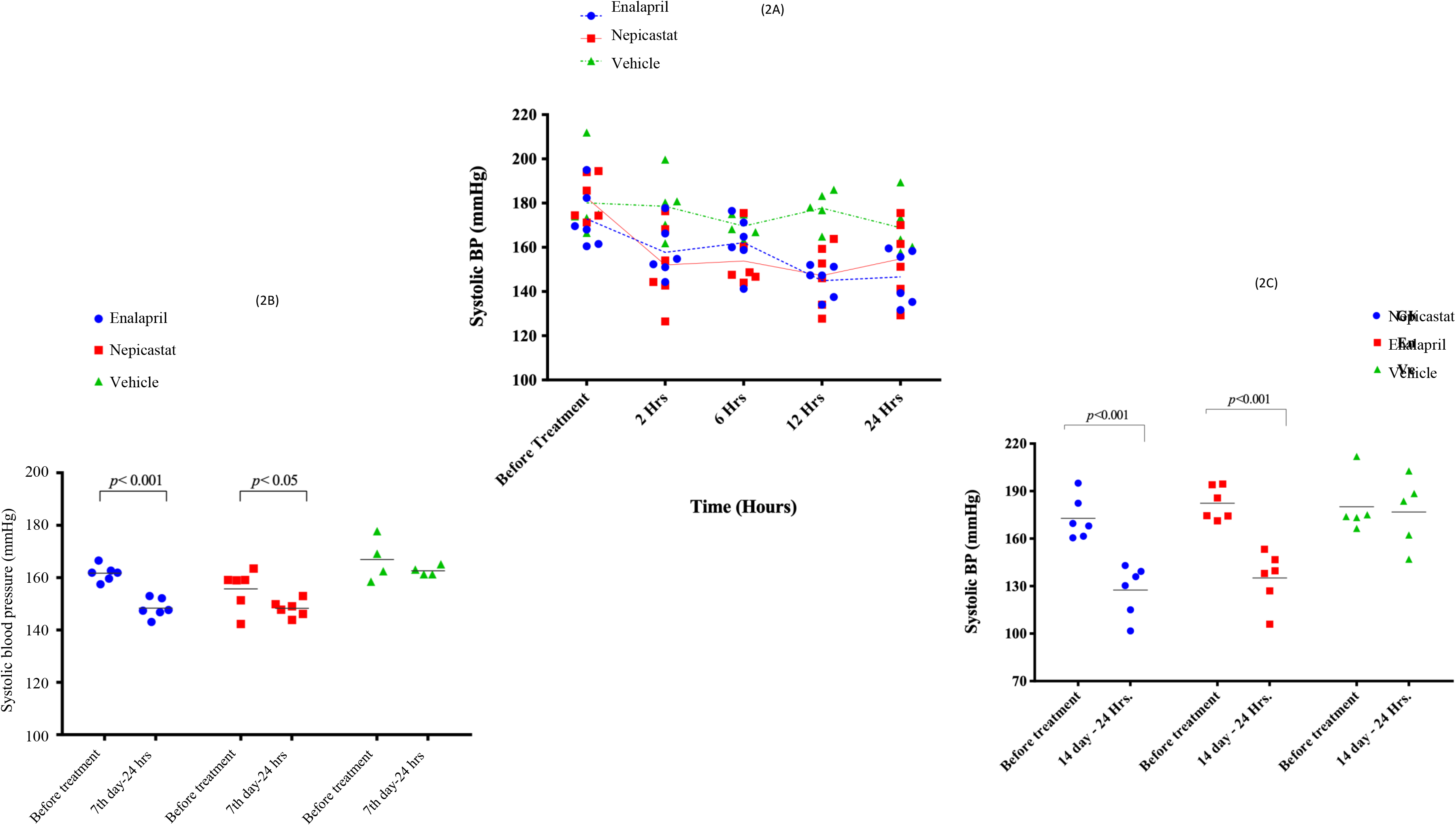

**Figure 3.**
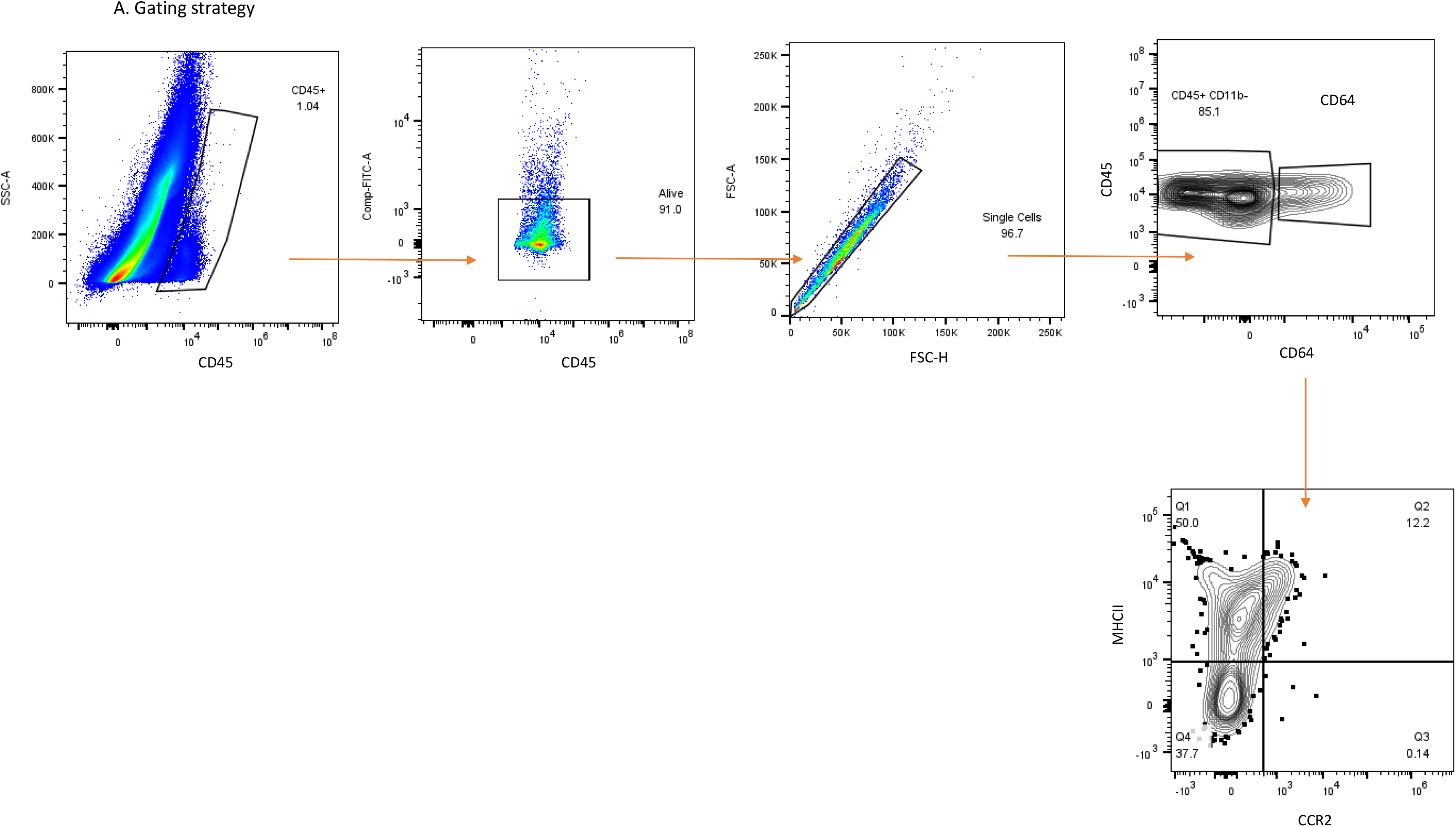
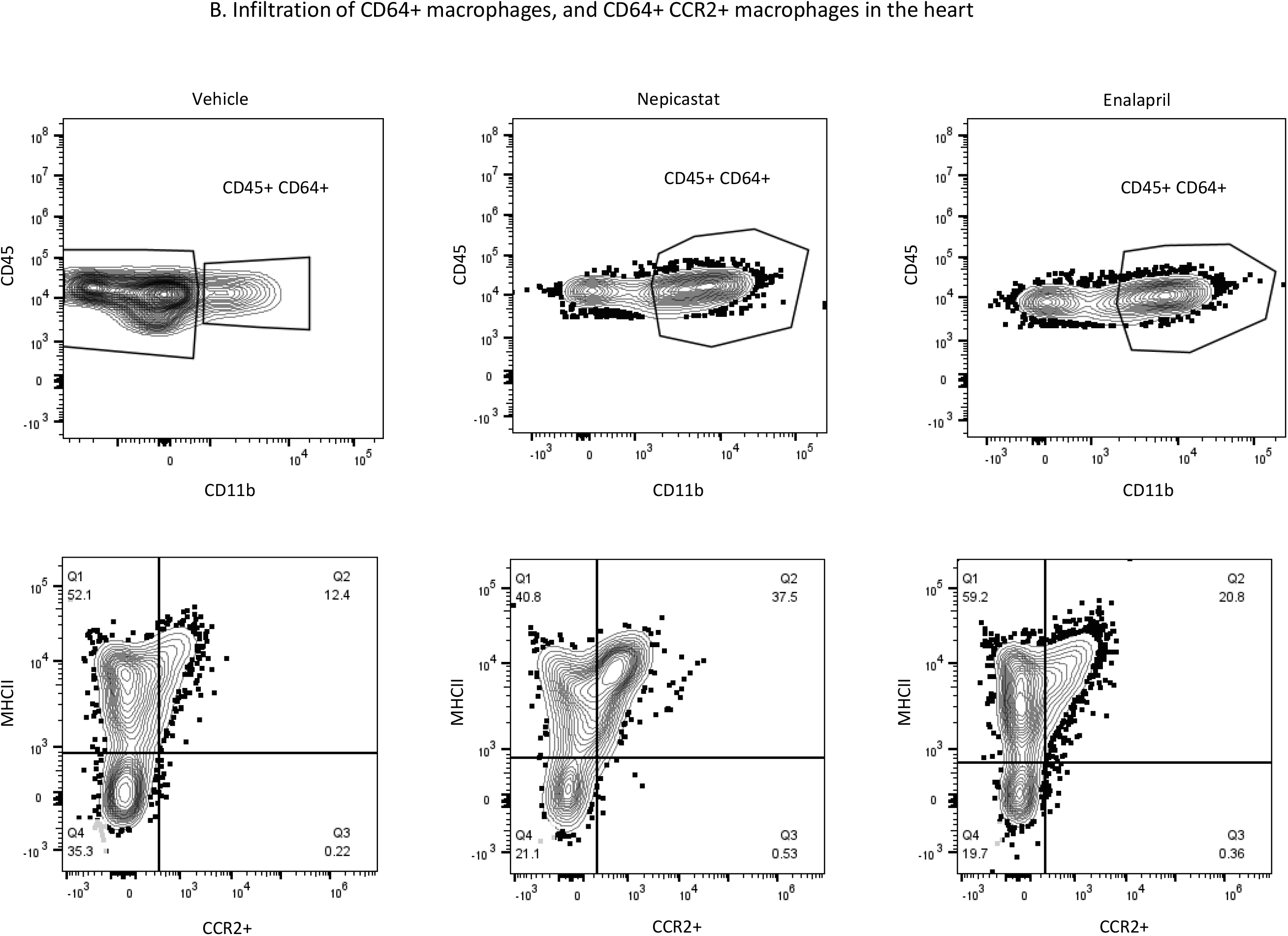

Flow cytometry analysis revealed infiltration of immune cells especially CD64+ macrophages and CD19+ B cells in the heart of nepicastat-administered SHR rats. We did not see a significant change in the CD4+ T cells or CD8+ cells in the heart of these rats. These results are consistent with the histopathology data.

## Discussion

Cardiovascular diseases especially hypertension is one of the major risk factors associated with the development of brain stroke. The current antihypertensive therapies pose several risk factors and side effects. To counter such off-target effects, we explored using acute and chronic studies if nepicastat which binds the novel target DBH offers alternatives to existing drugs with fewer side effects. We also investigated the toxicity or side effects of nepicastat administration in the SHR heart, lung, and kidney tissues. To our knowledge, it is a novel attempt to investigate if inhibiting the DBH enzyme could affect the toxicity/side effect in other organ systems, especially the inflammation/immune cells profile of the heart of nepicastat-administered rats. A repertoire of physiological, biochemical, and molecular biology methods was employed to characterize the effects of DBH inhibition in SHR rats, these observations were compared with enalapril or vehicle control groups. DBH inhibitors including nepicastat used in the past, were used as blood pressure-lowering therapies, none of these studies investigated the side effects or off-target effects on the heart or other tissue systems ^69^. This study is thus an advancement over the earlier ones and thus suggests the maximum blood pressure lowering timepoint. This study also was also provided evidence that neither the heart nor the kidney tissues were damaged nor did the rats suffer from abnormal behavior.

The systematic testing of DBH inhibitors can optimize the extensive drug screening and further their eventual translation as drugs. It is thus essential to test the DBH inhibitors in other relevant models like DBH gene knock-out or over-expressing models, which is beyond the scope of the present investigation. Additional molecular studies are also warranted, along with measurement of hemodynamic parameters. Nonetheless, our study has identified the important role of DBH inhibitor nepicastat and opened up new opportunities to investigate DBH as a novel druggable target for hypertension and cardiac hypertrophy.

## Materials and Methods

### Animal experiments

All animal experiments were carried out after obtaining approval from the Institutional Animal Ethics Committee (IAEC) of the Central University of Haryana (CUH), India (Protocol No. IAEC/570/CUH/2022). We strictly followed the guidelines of the Committee for the Purpose of Control and Supervision of Experiments on Animals (CPCSEA), Government of India. Three months old, eighteen male spontaneous hypertensive rats (SHR) were procured from CUH animal house, housed in groups, and maintained under the condition of 12 hours day/light cycle. After acclimatization for two weeks, animals were screened for hypertension by noninvasive blood pressure (NIBP), ECG, and echocardiography monitoring of rats.

### Pilot study

A pilot study (single dose study) was performed to check the antihypertensive potential of the nepicastat drug. Eighteen animals were randomized into three different groups (n=6 in each group), Control (Vehicle/5 % DMSO, 1ml/kg, ip.), Enalapril (10 mg/kg, orally) and nepicastat (30 mg/kg, ip.). After administration of a single dose of vehicle, enalapril, and nepicastat to SHR, systolic blood pressure was recorded at 2 hrs., 6 hrs., 12 hrs., and 24 hrs. using the NIBP system.

### Acute study

A 5-day acute experiment was performed in animals. Vehicle (5% DMSO, ip, daily), enalapril (10 mg/kg, orally, daily), and nepicastat (30 mg/kg, ip, daily) were administered to rats (n=6 in each group) continuously for 5 days. Twenty hours after drug administration, systolic blood pressure was monitored in all the animals.

### Chronic study

A 10-day chronic experiment was performed in animals. Vehicle (5% DMSO, ip, daily), enalapril (10 mg/kg, orally, daily), and nepicastat (30 mg/kg, ip, daily) were administered to rats (n=6 in each group) continuously for 10 days. Blood pressure was regularly monitored. On the 10th day, 6 and 24 hours after drug administration, systolic blood pressure was monitored in all the animals. Electrocardiography (ECG) and echocardiography parameters were also recorded from these animals at 24 hours after drug administration. Urine and blood samples were collected for molecular analysis before sacrificing the animals.

### Non-invasive blood pressure (NIBP)

Animals were restrained in a rat restrainer and allowed to acclimatize for 15 minutes. After the stabilization of the pulse, systolic blood pressure (SBP) was recorded using the NIBP system (IITC Life Science Inc., CA). Three or more blood pressure readings were taken from each animal and an average of three readings was considered as final SBP.

### Electrocardiography (ECG)

Animals were anesthetized using isoflurane anesthesia (5% Isoflurane mixed with 100% O2). After the stabilisation of the heart rate, ECG was recorded. Three measurements were performed from each animal and an average of three readings was considered as the final ECG parameter.

### Echocardiography

Animals were anesthetized using a combination of ketamine (50 mg/kg) +xylazine (5 mg/kg) anesthesia. The thoracic region of the animal was shaved and left ventricular hypertrophy was assessed using 2-D color Doppler echocardiography (Philips). Left ventricular physiology and function parameters such as systolic left ventricular intraventricular septal thickness (IVSs), diastolic left ventricular intraventricular septal thickness (IVSd), systolic left ventricular internal dimension (LVIDs), diastolic left ventricular internal dimension (LVIDd), systolic left ventricular posterior wall thickness (LVPWs), diastolic left ventricular posterior wall thickness (LVPWd), left ventricular fraction shortening (LVFS) and left ventricular ejection fraction (LVEF) were recorded. Three measurements were performed from each animal and an average of three readings was considered as the final echo parameter.

### Urine collection

Animals were kept in metabolic cages until 24 hrs. after drug administration and urine was collected. Urine immediately after the collection was stored at -20 °C until analysis.

### Plasma and tissue collection

Blood samples were collected before the sacrifice of animals and plasma was isolated from blood as per standard protocol. After sacrificing the animals, heart, lung, liver, and kidney tissues were collected for histology and molecular analysis. Tissues were stored in 10 % neutral buffered formalin for histology and in RNA later for molecular analysis.

### Histology

After sacrificing the animals, heart, lung, liver and kidney tissue cross sections were collected in 10% neutral buffered formalin (fixative). After 24 Hrs., cross sections were paraffin-embedded using the standard protocol. 5 μm sections were then taken on slides by cutting the paraffin blocks using microtome (Leica) and stained for Hematoxylin & Eosin and Trichrome ((HT10516-500ML, SIGMA ALDRICH, USA). The stained sections were analysed using Nikon Eclipse 55i microscope (20X) for myocyte, vascular changes, and collagen deposition respectively.

## Acknowledgments

We thank the University Grants Commission, Government of India for funding this study.

